# Simulating 3D refractive index distributions of suspended cells

**DOI:** 10.1101/2024.10.14.618263

**Authors:** Francesca Borrelli, Daniele Pirone, Lisa Miccio, Pasquale Memmolo, Pietro Ferraro, Vittorio Bianco

**Affiliations:** CNR-ISASI, Institute of Applied Sciences and Intelligent Systems “Eduardo Caianiello”, Via Campi Flegrei 34,80078 Pozzuoli (Napoli), Italy

**Keywords:** phase contrast tomography, in-flow holographic tomography, 3D refractive index, numerical simulation, cell phantom

## Abstract

The introduction of holo-tomographic flow cytometry has unlocked the possibility of imaging the 3D refractive index (RI) distribution of suspended cells while they flow and rotate in microfluidic streams. Similarly, approaches that optically trap and rotate the samples can image them in suspension conditions. Great effort has been spent in developing robust algorithms for the tomogram estimation, as well as denoising and 3D segmentation algorithms. However, the lack of a ground-truth dataset for suspended cells significantly hinders the development of image processing pipelines, limiting the advancement of the associated technology field. Here we propose a novel method for simulating 3D refractive index tomograms of suspended cells. We start with prior knowledge of the statistics of the 3D RI distribution and morphometry of various cell sub-compartments gathered from holo-tomographic flow cytometry experiments and combined with literature data to create realistic 3D distributions. Next, we introduce a simple method to obtain a simulation of the tomogram obtainable from each cell, which approximates the effects of the limited numerical aperture of the system, the speckle noise and the inversion algorithm itself. As a benchmark for the simulator, we created a shared dataset of RI tomograms with various levels of complexity, simulating yeast eukaryotic cells at different budding stages with various phenotypes of cytoplasmic vacuoles, and the presence of cytoplasmic vacuoles and lipid droplets in monocyte cells. We believe that this shared dataset and the simulation method will contribute to the development of a wide plethora of tomogram estimation and processing algorithms.

## 1. Introduction

Since its introduction, Digital Holography (DH) has become an extremely valuable Quantitative Phase Imaging (QPI) technique thanks to the unique advantages it offers, compared to the other methods belonging to the same class [1]. DH allows the visualization, under the weak scattering assumption, of phase samples such as cells or biological tissues. The acquired digital holograms of the specimens probed in transmission mode contain the whole information about the complex field diffracted by the specimen. By numerically processing the hologram, Quantitative Phase Maps (QPMs) are extracted, which are measurements of the optical path delay of the light passing through the sample, which makes DH a quantitative technique. DH offers the possibility to image in-vivo samples, without the need for chemical stains or fluorescent dies, since the phase information is retrievable from the hologram encoding; another huge benefit is given by the possibility of refocusing the object in post processing, which boosts the throughput of the acquisition since samples in continuous flow can be analyzed. A further step in the imaging of biological samples can be obtained by exploiting holographic tomography techniques [2]. QPMs contain an integral, coupled information about the Refractive Index (RI) of the sample and its physical thickness. Through tomography, it is possible to decouple this information by acquiring QPMs relative to different orientations between the sample and the probing light. With a proper reconstruction algorithm, the 3D RI map of the sample can be retrieved, which represents the most complete information about the sample of interest. There are two ways of obtaining the relative movement between the sample and the probing light. The first, most intuitive way, is to stabilize the position of the sample though opportune supports and scan the probing light around it [2]. In accordance, this method is referred to as illumination rotation paradigm. This method offers great advantages such as the precision of the illumination and the a-priori knowledge of the projection angles which can be finely controlled, reaching also sub-grade spacings. However, this technique also leads to great disadvantages such as the reduced throughput, due to the need of physically placing the sample on the support, and the mechanical stress this procedure induces over the sample that can get deformed as a result. In addition, due to a non-complete coverage of the illumination angular range, this methodic leads to the missing cone problem [3, 4], which results in severe artifacts in the reconstruction. The second relative rotation approach consists of fixing the position of the scanning light and moving the sample, in a paradigm referred to as sample rotation [2]. This possibility has recently been demonstrated for DH in a technique referred to as holo-tomographic flow cytometry [2]. The sample rotation is obtained as it flows into a microfluidic channel by creating a gradient in the flow speed. This technique strongly improves the overall throughput of the analysis and allows a more complete angular coverage of the sample but has the strong drawback of not knowing a priori the projection angles, which must be numerically recovered. Since both the rotation paradigms discussed above are prone to numerical errors and the presence of speckle noise, it is of crucial importance to possess numerical models to be used as ground truth for the development of robust tomographic solvers, denoising, image enhancement, and 3D segmentation algorithms. The problem of lack of universal ground truths has been long known in microscopic imaging, as it is often impossible to have access to the true morphology of the samples of interest [5]. Therefore, often the results have to be cross-validated using different imaging techniques or statistics to prove the correctness of the obtained reconstructions [6]. In the case of QPI, the verification of experimental results was traditionally made exploiting very simple objects such as uniform micro spheres [7, 8, 9] or simple numerical models [10, 11, 12], until more sophisticated methods have been proposed to print a test target through photolithography [13, 14, 15]. While this method is perfectly suitable for illumination scan rotation approaches, it is more difficult to conjugate it with in-flow tomography based on the rotation of suspended cells, which still strongly relies on numerical phantoms [16]. Great attention must be therefore put into the design of a reliable numerical phantom, able to represent the specimen in the most reliable way and tackle the reconstruction strengths and drawbacks of different solvers, taking into account also the optical properties of the chosen imaging system [17]. In this paper, we propose a numerical phantom simulation suitable for different cell lines at different levels of complexity. The statistics of these simulations were studied and extracted from the literature and from data measured by the holo-tomographic flow cytometry system on suspended cells. In particular, we apply the method by simulating the RI tomogram of a yeast cell with cytoplasmic vacuoles inside, based on the studies reported in [18], of a monocyte with lipid droplets (LDs) based on the findings reported in [19], and of a monocyte with vacuoles. We chose to benchmark the method using these test cases due to the availability of prior information from experimental tests carried out by our group and, most importantly, due to their significance for basic research in biology and diagnostics. Yeast cells are important eukaryotic cells used as a model for the study of cytoplasmic vacuoles. At any stage of their life cycle, they are characterized by the presence of a large roundish vacuole along with the potential presence of much smaller vacuoles with low RI values. In mammalian cells, e.g. in monocyte cells of the bloodstream, the presence of various phenotypes of cytoplasmic vacuoles (that can vary largely in number and size) is considered as a biomarker for several disfunctions and pathological conditions, e.g. in lysosomal storage diseases (LSDs) [20], SARS-CoV-2 [21] and other viral infections [22-24], and cancer [25-29] to name a few. Instead, LDs are intracellular compartments mainly located in the cytoplasm [30]. They were initially recognized only as storage organelles, but it has been discovered that they play multiple roles in maintaining intracellular homeostasis and that they dynamically interact with various organelles, such as mitochondria and lysosomes, thus affecting cellular processes [31]. For this reason, the involvement of LDs has been recognized in several pathologies, such as cancer [32], diabetes [33], or neurodegenerative diseases [34]. In particular in immune cells (e.g., monocytes) LDs are markers of viral infections [35] and inflammations [36], as their number and size increase in response to these stimuli. Once the 3D RI distribution of the cells is simulated, a simplified ad-hoc method is proposed to simulate the effects of the optical system and non-ideal inversion algorithms. Thus, the simulator generates realistic representations of the 3D RI tomograms acquired by noisy optical systems with limited numerical aperture and generated by inversion algorithms that introduce numerical artefacts. The whole method is then applicable to a larger variety of cells and organelles, provided that prior statistical information are available on the morphometry and RI of the sub-compartments to be simulated. We share the simulated dataset with the aim of providing a tool for the development of phase-contrast tomography algorithms in the absence of a ground-truth.

## 2. Numerical simulation of a phantom cell

### 2.1 General cell simulation

We developed a versatile algorithm for generating numerical phantoms of suspended cells, which we then tailored to simulate the unique characteristics of various cell lines, including yeast cells and monocytes with LDs or cytoplasmic vacuoles. Accordingly, four classes of sub-cellular compartments were modelled, i.e. external membrane, cytoplasm, nucleus and vacuoles for yeast cells and monocytes, or LDs for monocytes. Concerning the dimension and the shapes of the overall cell and of each intracellular structure, in order to obtain a model as realistic as possible, these parameters were varied for each realization and drawn from suitable distributions modelling real samples. In the simulation of the monocyte, the objective was to model as accurately as possible the LDs as studied in [19]. Therefore, even though the shapes of the overall cells, the membrane, the cytoplasm and the nucleus were changed in each simulation to ensure intervariability, their RI values were left constant, in particular RI_membrane_= 1.350, RI_cytoplasm_ = 1.365, RI_nucleus_ = 1.380. Differently, concerning the modelling of the yeast cells, a finer simulation of their sub-compartments of interest was realized. In [18], for the first time each structure of yeast cells at different life stages was studied and characterized through a specificity-enabling method, providing valuable information that are exploited here to create an accurate model. Accordingly, not only the shapes of the substructures are simulated based on [18], but also the changes in RI values for each stage of division of the budding yeast.

As a first simulative step, a box region of 200×200×200 pixels, with pixel size 0.125 μm, created, where the cell will be inserted. The external shape of the cell is simulated by drawing its volume and sphericity index from a realistic distribution inspired by [37, 38] and adapted to the physical dimension of the cells of interest. Concerning the yeast cells, they occupy around 1/3 of the modelled volumetric box region. Monocytes are bigger and occupy around 60% of the simulated volume. Once those values are calculated, the cell is as first order modeled as an ellipsoid whose semiaxes in a cartesian coordinate system are labelled as r_x_, r_y_ and r_z_. The r_x_ value is drawn from a gaussian distribution describing the mean distance of the cell center from the membrane, while r_y_ and r_z_ are univocally determined by considering the volume of the cell, the sphericity index and imposing the relation r_x_ < r_z_ < r_y_. Then, the ellipsoid modelling the cell is numerically altered to obtain an irregular, more realistic external shape. Subsequently, the cytoplasm region is modeled in a similar manner after drawing its three semiaxes values from opportune gaussian distributions, taking into account the volumetric proportion between cytoplasm and the overall cell. Setting the cytoplasmatic semiaxes to a smaller value compared to the ones of the overall external cell allows obtaining a leftover border modelling the external membrane. Concerning the nucleus, its center is randomly placed into the cytoplasmic region and varied over the different realizations of the simulated dataset. Its dimension is determined by first choosing its sphericity index, values of minor and major principal axes, and nucleus to cell ratio in accordance with the known values. The latter parameter is particularly important for yeast cells as they are characterized by important changes in nucleus dimensions depending on the cell life stage. Consequently, the volume of the nucleus is calculated in accordance with the volume of the cell through the nucleus to cell ratio. The third principal axis is then univocally determined. Once the geometries of the mentioned structures have been determined (external membrane, cytoplasm, nucleus), their RI values have to be modeled.

### 2.1.2 Simulation of cytoplasmic vacuoles

Particular attention was set into the modelling of the vacuoles, as they represent the most distinctive trait of eukaryotic yeast cells and have been linked to different pathological conditions in mammalian cells, e.g. their presence, number and typical patterns are bioindicators of the presence of inflammatory diseases, stress conditions, virus induced alterations or lysosomal storage diseases (LSD), as mentioned above. Moreover, it is possible to experimentally control the volume and the number of these organelles, engineering the yeast cells acting on the osmotic conditions of the surrounding buffer, or in the case of mammalian cells by using external drugs to simulate this phenotype. Hence, it is important to implement a simulation tool capable of describing all these conditions. Here, each vacuole is modeled as an ellipsoid whose center is randomly allocated into the cell, respecting the experimentally observed statistics of vacuole placement. Indeed, depending on the cell stage and on their sizes, vacuoles tend to occupy different positions in the cytoplasm of the cell. Then, the sphericity index of the vacuole is drawn from the statistics in [18]. The most important factor used for the parametrization of the vacuole is its equivalent diameter. By selecting this value and combining it with the sphericity index, the three semiaxis of the ellipsoid vacuole r_xv_, r_yv_ and r_zv_ are calculated.

As described in [18], the vacuole region is characterized by a higher value of RI in its membrane region, which we refer to as perivacuolar region, and then by a value of RI gradually decreasing up to the center of the vacuole. To model this, we first randomly generate the value of RI of the perivacuolar region RI_perivac_ and of the central vacuolar region RI_vac_ considering the reported statistics. Then, the internal area is modelled through concentric ellipsoidal shells. Once the number of shells, *N*, is selected, depending on the equivalent radius of the vacuole, their geometry is described as reported below.

- The most external shell (n = 1) coincides with the generated ellipsoid representing the entire vacuole. In formulas, r_x1_ = r_xv_, r_y1_ = r_yv_ and r_z1_ = r_zv_.
- The semiaxes of the most internal shell (n = N) are equal in size to the 10% of the ones of the most external shell. In formulas, r_xN_ = 10%r_xv_, r_yN_ = 10%r_yv_ and r_zN_ = 10%r_zv_.
- An equally spaced vector of shells’ semiaxes is generated for each dimension, ranging from the calculated outermost and the innermost ones, and then assigned to each shell. In formulas s_x_ = *linspace*(r_xN_, r_x1_, N), s_y_ = *linspace*(r_xN_, r_x1_, N), s_z_ = *linspace*(r_xN_, r_x1_, N), where s_x_ is the vector containing the x semiaxes of each shell, s_y_ the vector containing y semiaxes and s_z_ the vector containing the z semiaxes. The MATLAB® function *linspace* returns a vector of N elements equally spaced from the first argument (r_xN_ in this case) to the latter (r_x1_ in this case).
- Similarly, an equally spaced distribution of mean RI values is created for each shell, ranging from the RI assigned to the peri-vacuolar region to the one assigned to the internal vacuole. In formulas, RI_mean_ = (RI_meanN,_…,RI_mean1_) = *linspace*(RI_perivac_, RI_vac_, N).
- A distribution with mean equal to 1 and standard deviation equal to 0.03 is used to generate a standard deviation value for each shell. In formulas, stds = (std_N_…std_1_).
- The final RI value for each shell n is drawn from a gaussian distribution with mean RI_mean,n_ and standard deviation std_n_.
- Finally, a smoothing operation is applied to the entire vacuolar region to smoothen the sharp variations between the shells followed by a volumetric erosion equal to the 22% of the value of the equivalent radius, to reduce the width of the peri-vacuolar region.

Fig. 1 sketches the geometrical structure of the proposed vacuole before the smoothing operation.

**Fig. 1:**
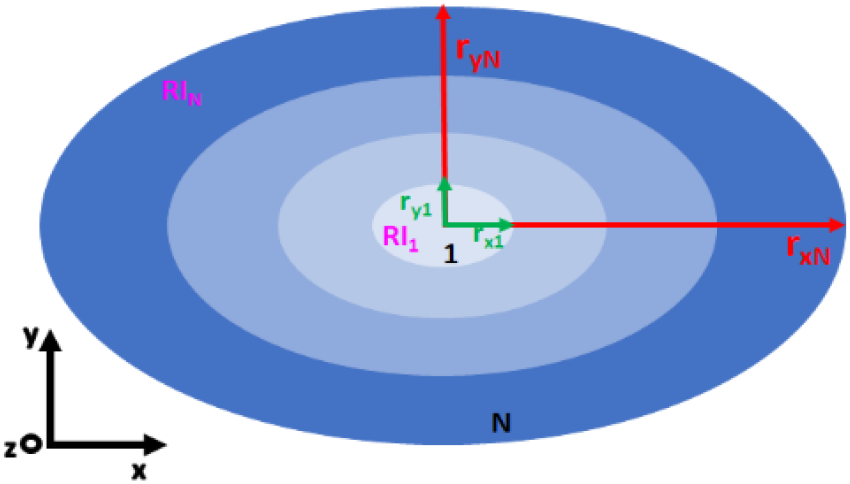
schematical representation of the geometric structure used for modelling the cytoplasmic vacuole.

### 2.2 Simulation of lipid droplets

LDs represent structures with high RI values linked to several regulatory mechanisms in living cells, both in physiological and pathological states. As for the case of the vacuoles, these organelles are modeled setting their equivalent radius as the principal parametric parameter. Like all the organelles discussed, LDs are modeled as ellipsoids. Combining the value of equivalent radius with the sphericity index and the volume, as discussed in [19], the three semiaxes are determined. The number of LDs simulated is chosen considering the ratio between the LDs aggregate volume and the volume of the overall cell. Knowing the average volume of each LD, the chosen number of LDs inserted in the cell is 4. As a last geometric consideration, it was noted in ref. [19] that the lipids tend to aggregate in a small area which is displaced from the centroid of the cell. Therefore, this specification is incorporated in the placement of the centroid of each LD. Concerning the assignation of RI, its mean value and standard deviation were reported in ref. [18]. However, it was proven that the RI profile is not constant in the LD volume but increases going towards the center. Accordingly, we exploited a simulation mechanism similar to the one used for the vacuoles, where three shells were exploited to model the RI profile:

- The most external shell coincides with the generated ellipsoid representing the entire LD.
- The semiaxes of the most internal shell are equal in size to the 30% of the ones of the most external shell.
- The semiaxes of the median shell are calculated as the mean value between the internal and the external.
- The RI value of the most internal shell is drawn from a gaussian distribution with mean RI equal to the one of the overall LD and standard deviation = 0.001.
- The RI value of the most external shell is equal to the 70% of the one of the inner shell.
- The RI value of the central shell is the mean value between the internal and the external ones.
- Finally, a smoothing operation is applied to the entire LD region to smoothen the sharp variations between the shells.

Differently from the vacuole simulation, a smoother profile of RI is simulated, as the presence of a sharper LD membrane was not detected in ref. [19], thus explaining the presence of only three shells and the absence of the erosion needed to simulate the peri-vacuolar membrane. Finally, the mean value of the LD’s RI profile is checked to ensure it falls into the expected ranges.

### 2.3 Simulation of monocytes with cytoplasmic vacuoles

This simulation constitutes the merge between the two cases presented before. This case is very important for two main driving reasons:

i. Monocytes like all the other cells of the blood stream naturally live in suspension. Thus, this class of cells is the one that benefits the most from the use of a holo-tomographic apparatus working in flow cytometry mode, for which the absence of a ground-truth is a non-negligible impairment.
ii. The presence of cytoplasmic vacuoles inside monocytes is associated with several conditions of inflammation and chronic diseases, as mentioned in the previous sections. Thus, there is a strong interest in phenotyping patterns of cytoplasmic vacuoles inside these cells, some of them challenging the optical resolution of holo-tomographic flow cytometry systems.

In the monocyte cell introduced in 2.2, cytoplasmic vacuoles are modeled as presented in 2.1. In this case, small vacuoles are simulated, in a numerosity ranging from 7 to 10 for each cell. The equivalent radius of the vacuole was drawn from a gaussian distribution with mean 0.8 and standard deviation 0.8, since those structures are small compared to the overall volume of the cell. Concerning their mean RI value, it was drawn from a gaussian distribution with mean 1.360 and standard deviation 0.008.

## 3. Results

In Fig. 2, the simulation for two different yeast cells is reported. As it can be appreciated by the scalebar, yeast cells are small with respect to the majority of mammalian cells. In Fig. 2 (a)-(d), a yeast cell with one single vacuole of moderate dimensions (i.e. with a volume occupying approximately 10% of the cell’s volume) is depicted. The cell has a quasi-spherical external shell. In Fig. 2(a) and (c) the central slice (cuts of the simulated cell in the plane yz and xy respectively) of the simulated 3D RI distribution are reported. Visualization 1 shows the slice-by-slice scanning of the whole simulated 3D RI distribution. The corresponding isolevel representation is shown in Fig. 2(d). In Fig. 2(b) it is possible to visualize the cytoplasmic vacuole with the peri-vacuolar membrane, defined as in [18], whose structure can be observed in detail in the RI profile along the red dotted line shown in Fig. 2(a). The panels in Fig. 2 (e)-(h) report the simulation of a yeast cell characterized by a large roundish vacuole (i.e. with a volume occupying approximately 7.5% of the cell volume) and a multitude of smaller vacuoles (i.e. with a volume occupying approximately 0.3% of the cell volume each) in the peripheral region (Fig. 2(h)). The external shell of the cell is non symmetric. The difference in size and the internal structures can be appreciated from peripheral yz and xy slices of Fig. 2(g) and 2(e) respectively, and the corresponding RI profile along the red dashed line, shown in Fig. 2(f). Visualization 2 shows the slice-by-slice scanning of the simulated 3D RI distribution for this cell.

**Fig. 2:**
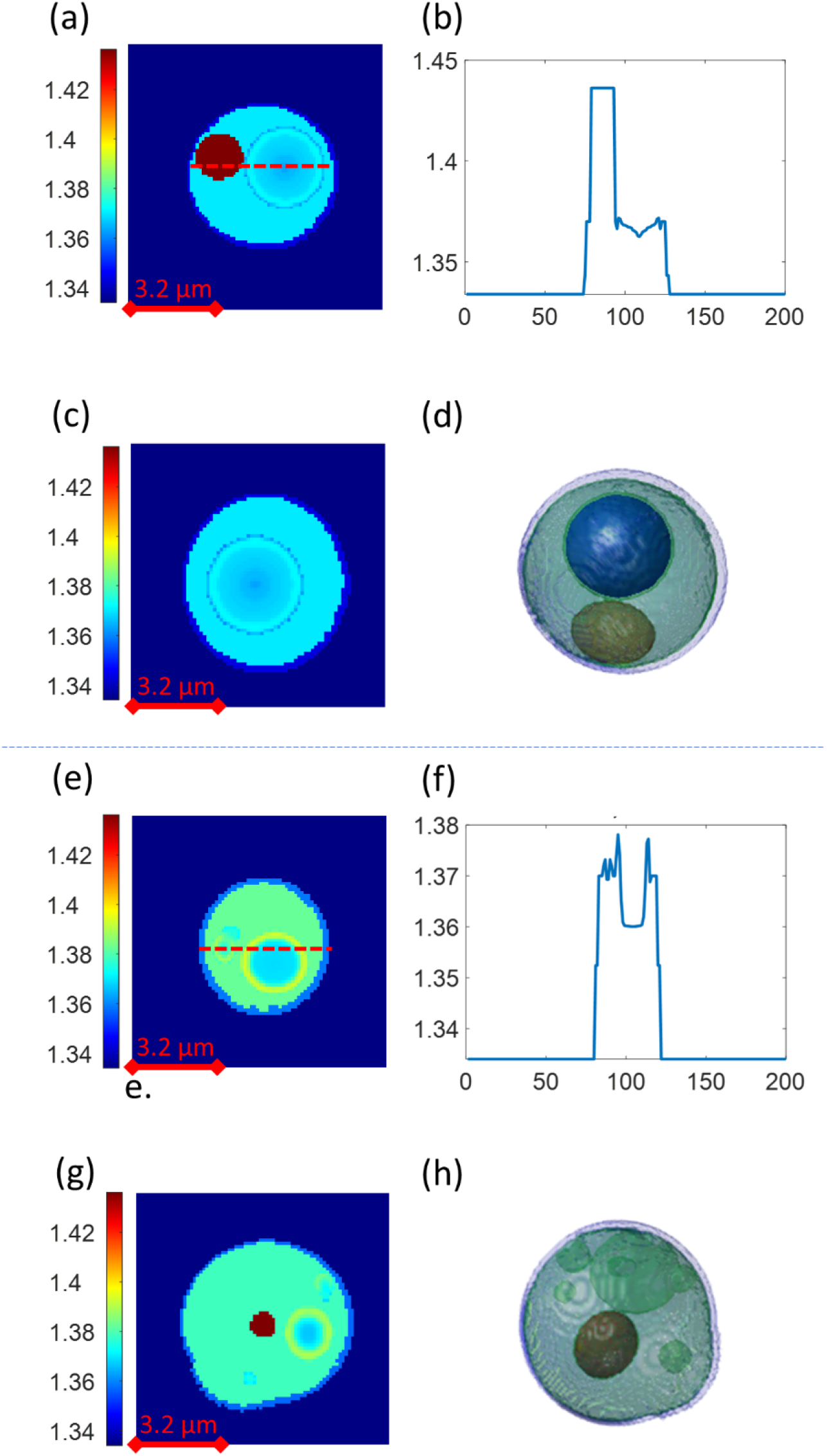
simulation results for (a-d) a yeast cell with a large vacuole inside (Visualization 1) and (e-h) a yeast cell with a large vacuole and a multitude of smaller vacuoles (Visualization 2). (a, g) Visualization of a slice in the yz plane from the 3D RI profiles; (b, f) Cut along the red dotted line in subfigures (a,e), highlighting the RI profile into the vacuole; (c, e) Visualization of a slice in the xy plane from the 3D RI profiles; (d,h) Isolevel visualizations.

Fig. 3(a), (b) reports the isolevel visualization of two different yeast cells in the late-G1 stage, according to the nomenclature reported in [18] that describes the different stages of the cell budding process. This simulation is aimed at showing the possibility of including the case of a cell that does not match exactly the common approximation of spheroidal external shell. Hence, eventual asymmetries in the external shell are simulated. Accordingly, the simulation is performed setting the simulation in agreement with the statistics reported for the set of experimental results obtained in the case of late-G1 stage [18]. An ensemble of 50 cells of late-G1 yeast cells is simulated and histograms of relevant statistics are computed. In particular, Fig. 3(c) shows the histograms relative to global parameters such as the overall volume of the cell and its equivalent diameter; the average value of RI and the value of the three principal axis of the cell. In addition, the nucleus’ volume, equivalent diameter and average RI histograms are reported, considering that, as mentioned in Section 2, the nucleus’ geometry is parametrized starting from the available value of sphericity index for a late-G1 cell. Finally, histograms of the vacuole’s sphericity index, average RI and average RI of the perivacuolar membrane are reported, considering in this case that the vacuole geometry is parametrized over the equivalent diameter, which is fixed for the late-G1 class of cells, thus providing a constant equivalent volume. As it can be inferred from the set of histograms, the simulated statistics, also summarized in Table 1, are in good accordance with the tables reported in [18], which confirms the reliability of the simulation.

**Table 1:**
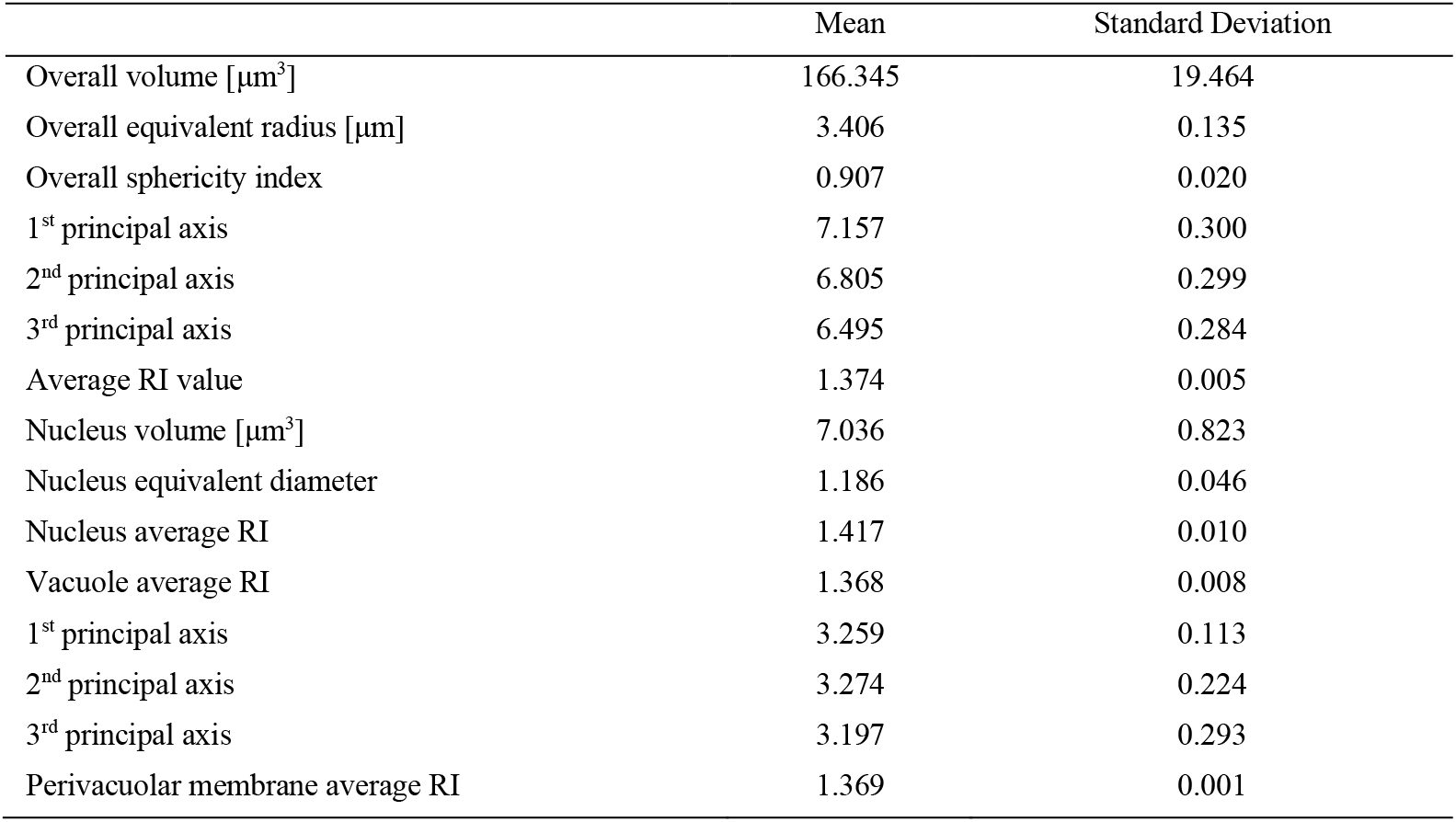
cell parameters measured over a population of 50 late-G1 simulated yeast cells.

|  | Mean | Standard Deviation |
| --- | --- | --- |
| Overall volume [$\mu\text{m}^3$] | 166.345 | 19.464 |
| Overall equivalent radius [$\mu\text{m}$] | 3.406 | 0.135 |
| Overall sphericity index | 0.907 | 0.020 |
| 1<sup>st</sup> principal axis | 7.157 | 0.300 |
| 2<sup>nd</sup> principal axis | 6.805 | 0.299 |
| 3<sup>rd</sup> principal axis | 6.495 | 0.284 |
| Average RI value | 1.374 | 0.005 |
| Nucleus volume [$\mu\text{m}^3$] | 7.036 | 0.823 |
| Nucleus equivalent diameter | 1.186 | 0.046 |
| Nucleus average RI | 1.417 | 0.010 |
| Vacuole average RI | 1.368 | 0.008 |
| 1<sup>st</sup> principal axis | 3.259 | 0.113 |
| 2<sup>nd</sup> principal axis | 3.274 | 0.224 |
| 3<sup>rd</sup> principal axis | 3.197 | 0.293 |
| Perivacuolar membrane average RI | 1.369 | 0.001 |

**Fig. 3:**
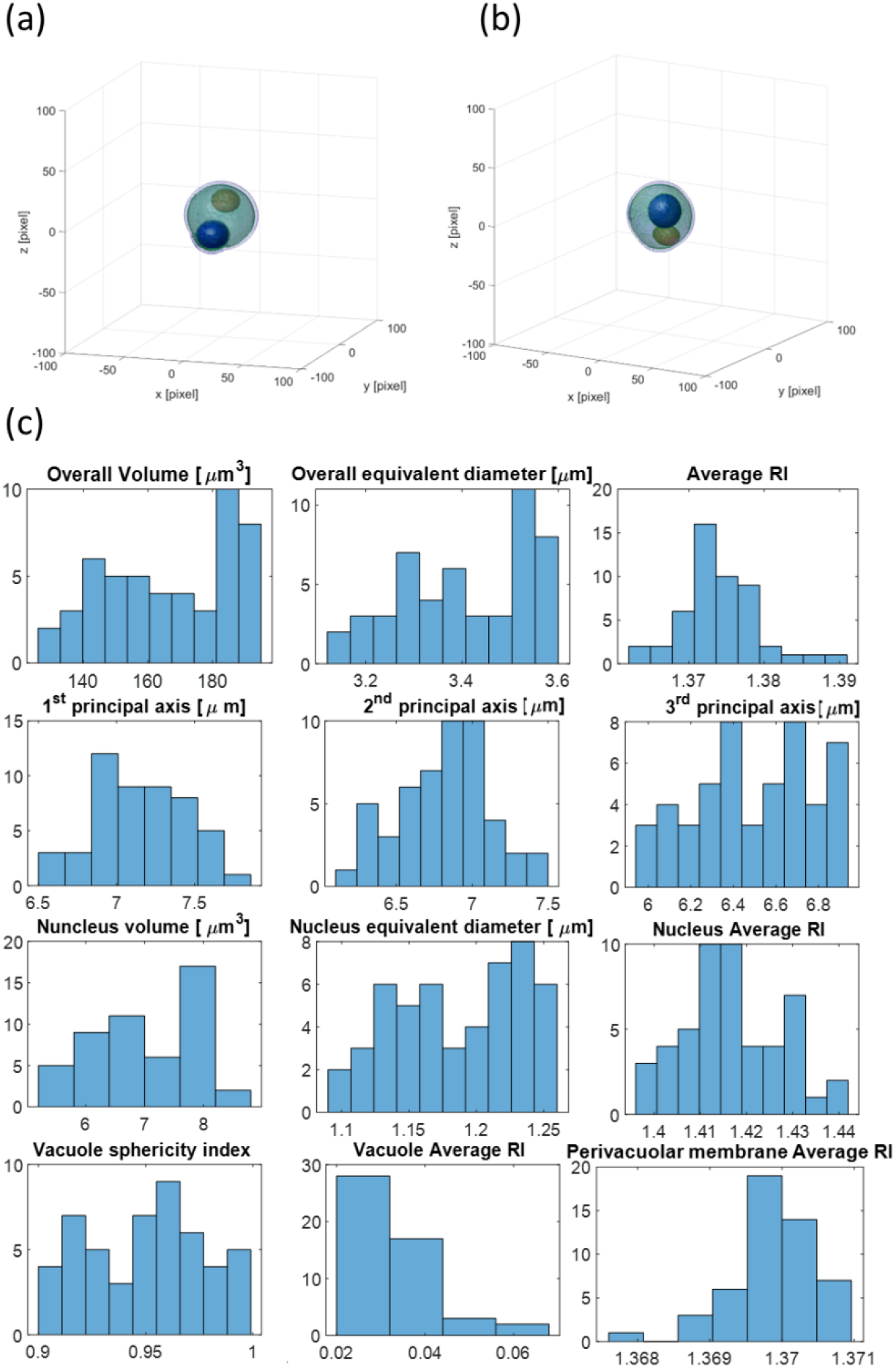
simulation results for late-G1 yeast cells. (a, b) isolevels of two different simulated cells; (c) Histograms of some relevant statistics evaluated over a population of 50 cells. The statistics are in good accordance with the ones in the literature.

Fig. 4 reports a similar analysis for the case of monocytes with LDs. In this case the simulation is performed setting the values for the external membrane, cytoplasm and nucleus as described in section 2.1 and the ones for the LDs as in [19], Table 1. As can be observed in the reported isolevel representation (Fig. 4(a), (b)), LDs tend to aggregate into a reduced portion of the cell’s volume, a behavior observed in [19] and that we included in our modelling. From this visualization, their volume can be quickly compared to the volume of the overall cell (each LD occupies about 0.006% of the whole cell volume). The slice-by-slice visualization of Fig. 4(b) (see also Visualization 3) allows appreciating the spatially variating RI distribution of each LD. Also in this case, statistics of relevant parameters relative to LDs are evaluated for a population of 50 simulated cells (sphericity index, average RI and RI standard deviation, reported in Fig. 4(c)) and show a good accordance with the values reported in [19].

**Fig. 4:**
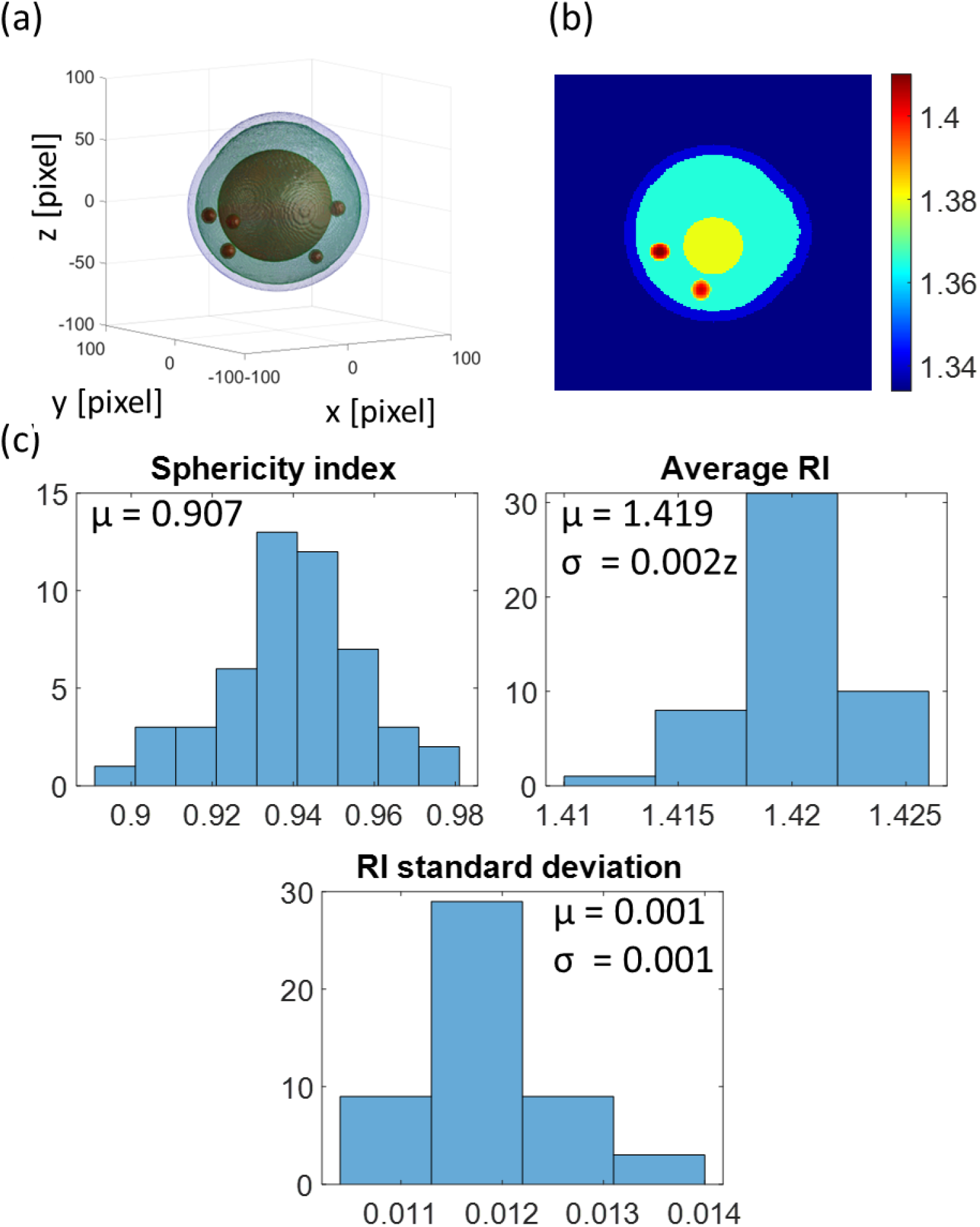
results for the simulations of a monocyte with LDs (Visualization 3). (a) Isolevel. From this visualization, the size and the positioning of the LDs can be appreciated, compared to the overall cell; (b) yz cut of the simulated 3D RI distribution. The non uniform distribution of RI into the LD can be appreciated; (c) Histograms of the sphericity index, average RI and standard deviation values are evaluated for the LDs in a population of 50 monocytes. μ = mean value. σ = standard deviation.

Finally, the results for the case of a monocyte with simulated cytoplasmic vacuoles are presented in Fig. 5. Fig. 5(a) presents an isolevel and a peripheral yz slice from the cell, where the asymmetry of the cell’s external shell can be noted, together with the placement of the vacuoles and of their non-uniform RI distribution. Visualization 4 shows the slice-by-slice scanning of the whole simulated 3D RI distribution of Fig. 5(a). In Fig. 5(b), (c), isolevel visualizations of two additional simulated cells are depicted.

**Fig. 5:**
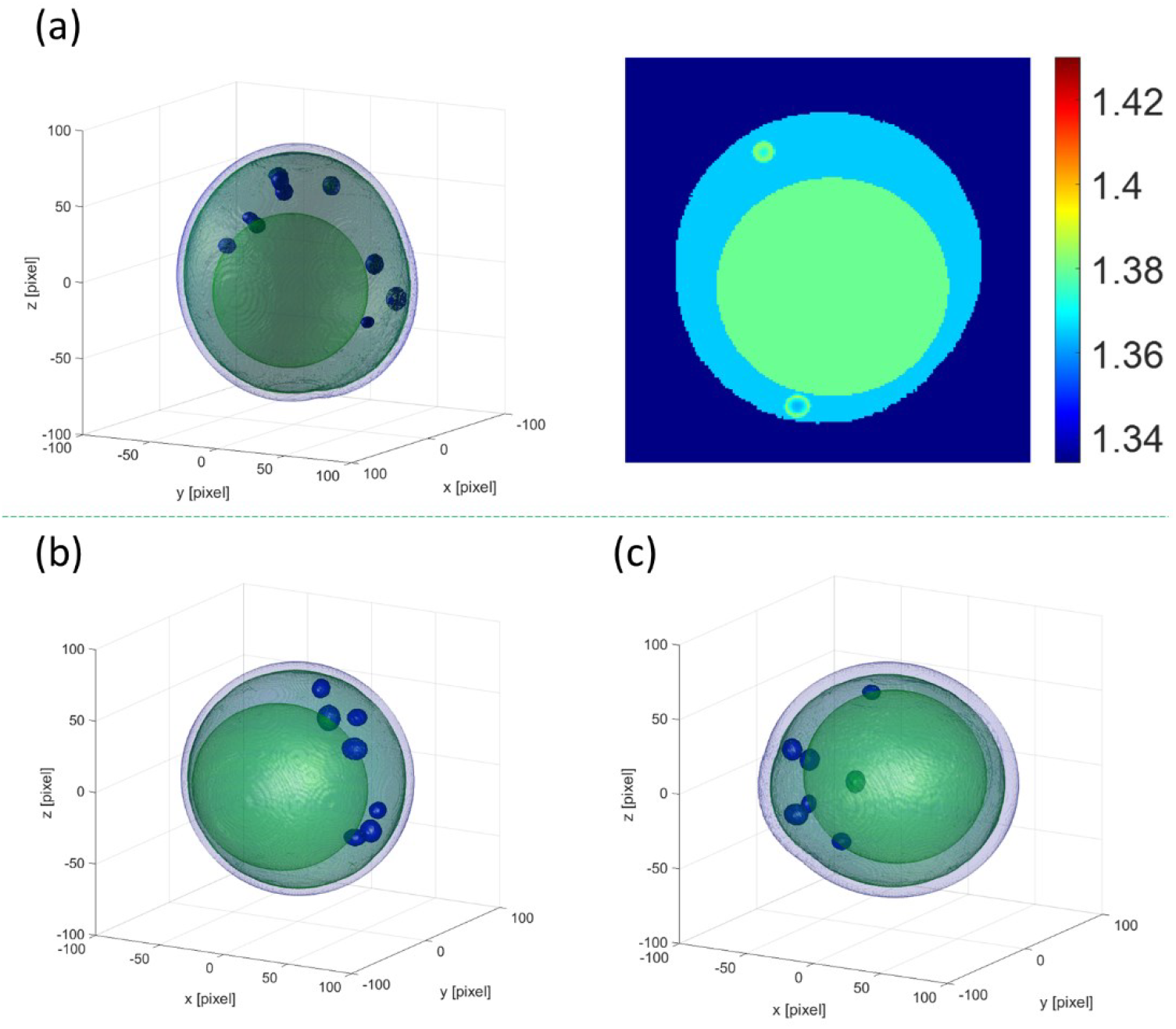
results for the simulations of monocytes with cytoplasmic vacuoles. Top panel, (a), an isolevel visualization and a yz slice from a simulated cell, highlighting the positioning of the vacuoles and their spatially-varying RI distribution (Visualization 4). Bottom panel, (b, c) isolevel of two other simulated cells.

### 3.1 From the simulated cell to its simulated reconstructed tomogram

After the ground-truth 3D RI distributions are simulated, the system Point Spread Function (PSF), speckle noise, and the influence of the employed inversion algorithm (e.g. Iradon) have to be considered. Here, a simplified general method is proposed, which allows to simulate and test each phase of the holo-tomographic acquisition-reconstruction pipeline. This is summarized in the sketch of Fig. 6(a). The starting point is the cell simulation we propose in this work. Convoluting this model with the Point Spread Function (PSF) of the optical system in use, the simulated tomogram model is obtained. Reprojecting this model along the available tomographic angles allows simulating the QPMs, which can be eventually corrupted by noise. Finally, composing these QPMs with a suitable tomographic solver allows computing the reconstructed, simulated tomogram. Fig. 6(b) represents each step of the presented pipeline applied to our in-flow tomographic flow cytometry optical setup for a sample simulated cell.

**Fig. 6:**
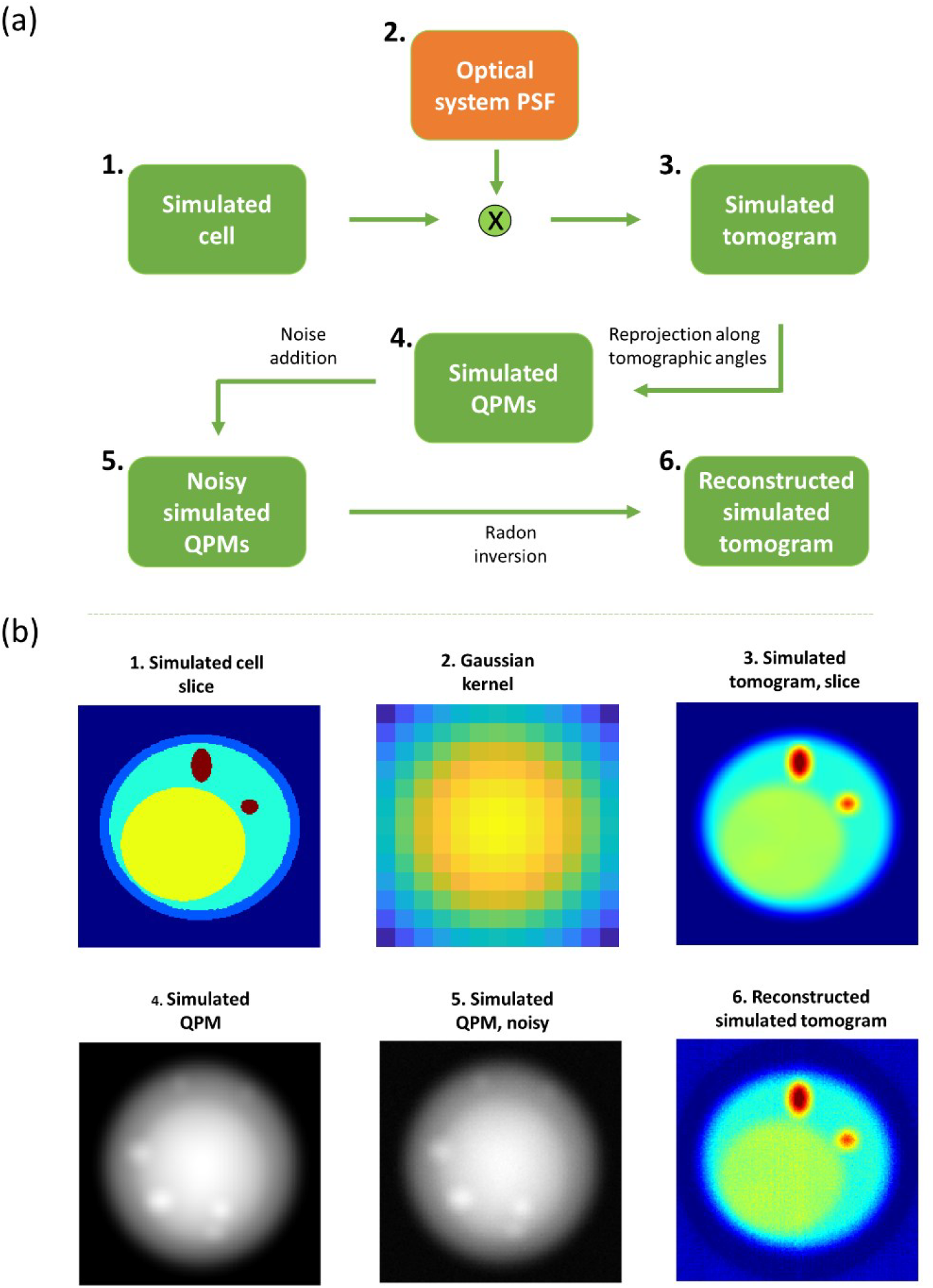
General sketch of a holo-tomographic acquisition-reconstruction pipeline. (a) Most important steps are schematically depicted; (b) Images corresponding to each step particularized to our optical setup and reconstruction pipeline are discussed (Visualization 5).

1. A RI distribution of interest, C, of size *n x n x n* is simulated according to the presented numerical models. Fig. 6(b1) sketches the central slice of C.
2. The PSF of the optical system is modeled through a 3D gaussian kernel G of size 13 *x* 13 *x* 13 and standard deviation 51. Fig. 6(b2) depicts the central slice of the gaussian kernel.
3. The simulated tomogram T of size *n x n x n* is obtained as the convolution between the modeled PSF and the RI distribution C. In formulas, *T* = *C*⨂*G*, where ⨂ is the convolution operator. Fig. 6(b3) sketches the central slice of T.
4. The QPMs are obtained by reprojecting T along the available projection angles. In formulas *Q* = *ψ*{*T, theta*}, where *ψ*{… } is the operator modelling the projection, *theta* is the set of projection angles with size 1 *x t* and *Q* is the stack of resulting QPMs of size *n x n x t*. In our case, the operator *ψ*{… } is obtained through the MATLAB® *radon* function. Fig. 6(b4) depicts one of the obtained QPMs.
5. The noisy QPMs, QPM_noise_, are obtained by adding a suitable noise distribution *N* over the uncorrupted QPMs. In our case, the noise is modeled as a gaussian distribution of size *n x n* with mean 0 and standard deviation 0.05. In formulas, *QPM*_*noise*_ = *QPM* + *N*. Fig. 6(b5) depicts one of the noisy QPMs.
6. The simulated reconstructed tomogram *T*_*rec*_ is obtained by inverting the operator *ψ*{… }, composing all the available projections and the associate projections angles with a suitable physical criterion. In our case, this step is performed resorting to the MATLAB® *iradon* routine. In formulas *T*_*rec*_ = *ψ*^−1^ {*QPM*_*noise*_, *theta*}. Summing up, the simulated reconstructed tomogram is obtainable from the simulated cell as:

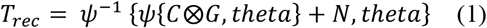

Fig. 6(b6) depicts the central slice of the simulated reconstructed tomogram *T*_*rec*_. Visualization 5 shows, in the slice-by-slice scanning mode, the side-by-side comparison between *C, T*, and *T*_*rec*_.

Fig. 7 depicts the application of this pipeline to the cells simulated and discussed in this work. The top row shows an yz slice from the simulated cells in Fig. 2, Fig. 4 and Fig. 5(a), while the bottom row shows the resulting simulated tomogram reconstruction obtained as described above. Each step of the pipeline is important and can be flexibly used to tailor suitable algorithms of interest according to the specific needs of the user and the features of the employed optical system. For example, we can investigate the effects of limited projection angles and the impact of errors in projection angle recovery, particularly in in-flow setups. In addition, denoising procedures can be experimented to study their effects on QPMs or even on the tomogram. Finally, tomographic solvers can be studied and fine-tuned, to compare the ideal tomogram to the reconstructed one.

**Fig. 7:**
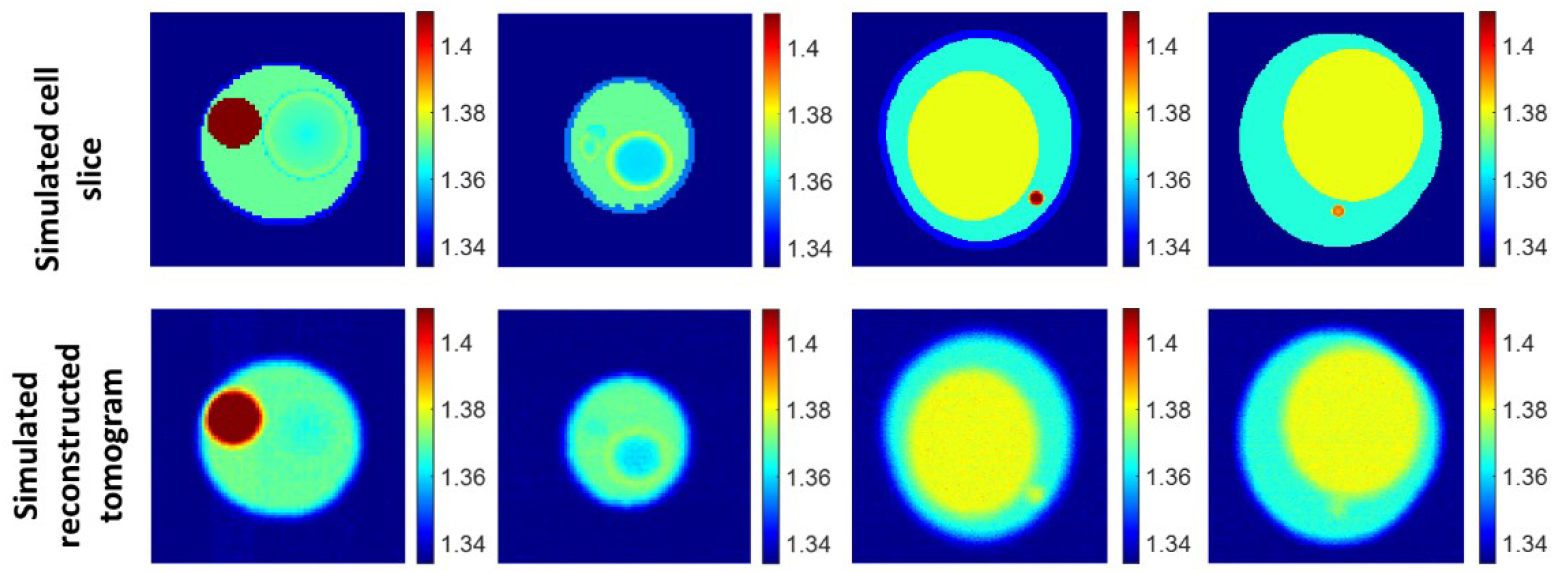
application of the introduced pipeline to the simulated cells displayed in this work. Top row, yz slice from the simulated cells; Bottom row, corresponding slice of the reconstructed simulated tomogram.

## 4. Conclusions

In this paper, a simulation tool has been proposed for the high-fidelity modelling of the 3D RI tomograms of suspended cells. As a benchmark, we simulated yeast cells as an example model for eukaryotic cell lines and human monocytes as an example model for mammalian cell lines. We simulated the 3D RI distribution first, accounting for the presence of internal organelles/sub-compartments. Thus, we created a dataset of cells with various levels of complexity. As a proof of concept, we simulated the presence of various phenotypes of cytoplasmic vacuoles inside yeast cells and monocytes, and the presents of LDs inside monocytes. All these compartments are of great interest since they are biomarkers associated with various dysfunctions, inflammations, stress conditions and diseases in cells of the human bloodstream. Besides the reported examples, the proposed approach is general enough to simulate realistic RI distributions of various organelle complexes provided that prior statistical information are available from the literature and/or measured tomographic data. Here, the simulations are based on experimental statistics present in the literature and in some cases on data gathered by using a holo-tomographic flow cytometry apparatus. Once the RI distributions of the cells are simulated, a simple method is proposed to obtain a realistic simulation of their estimated tomogram through a limited Numerical Aperture (NA) noisy system. In this way, the loss of resolution due to the passage through the system PSF, the tomogram quality worsening in terms of SNR due to the presence of speckle noise, and the numerical artefacts due to the non-ideal inversion algorithm (e.g. the Inverse Radon Transform) can be considered. Results show the versatility of the simulation method for the realization of numerical phantoms which are of primary importance for the development of holo-tomographic flow cytometry reconstruction, denoising, 3D segmentation and analysis algorithms in the absence of a ground-truth.

## Supporting information

Visualization 1

Visualization 2

Visualization 3

Visualization 4

Visualization 5

## Acknowledgements

This work was supported by a project PRIN 2022 – Label-free cytoplasmic vacUoles pheNotyping plAykit (LUNA) Prot. 960, 30th June 2023 – funded by the Italian Ministry of University & Research in the framework of the European Union program NextGenerationEU (2022BKYM22 - Project CUP: B53D23002490006).

## Data availability statement

The whole simulated dataset is shared publicly and can be downloaded at the following link: https://doi.org/10.6084/m9.figshare.27226464

## Conflict of interest statement

Authors declare no conflict of interest associated to this work.

## Notes

https://doi.org/10.6084/m9.figshare.27226464

### Competing Interest Statement

The authors have declared no competing interest.

